# Integrated RNA sequencing reanalysis reveals reproducible matrix-immune signatures in idiopathic pulmonary fibrosis

**DOI:** 10.64898/2026.06.24.734263

**Authors:** Sneha Nandimandalam, Jin He, George I. Mias

**Author notes:** **Author for correspondence:** George I. Mias.

## Abstract

In idiopathic pulmonary fibrosis (IPF), the lung is remodeled through coordinated epithelial, stromal, and immune-associated programs, but individual transcriptomic cohorts are often too small to separate shared disease signals from demographic and study-level variation. To increase statistical power while preserving study-aware interpretation, we integrated raw bulk lung RNA sequencing (RNA-seq) data from five well-annotated studies and analyzed 223 samples in a common framework that modeled sex, age, library layout and repeated sampling. IPF showed a broad and reproducible expression shift, with 2,443 genes meeting the differential-expression threshold of false discovery rate (FDR) *<* 0.05 and absolute log_2_ fold change at least 1. The dominant program combined extracellular matrix remodeling, stromal and epithelial activation, complement and B-cell-related pathways, cilium-associated processes, and relative depletion of oxidative phosphorylation and proteasome pathways. Sex-stratified analyses recovered a shared fibrotic core with smaller sex-skewed components, whereas age-related disease effects were weaker and centered on immune activation. A leave-one-study-out elastic-net analysis using fixed disease-gene panels classified IPF across held-out studies, supporting cross-study portability of the core signature. This integrated reanalysis strengthens evidence for a stable matrix-immune IPF program and reinforces the view that core disease-associated transcriptional programs are reproducible across heterogeneous cohorts.

## 1. Introduction

Idiopathic pulmonary fibrosis (IPF) is typically diagnosed in older adults, with fibrotic remodeling of the lung accompanied by declining respiratory function and poor survival [1,2]. Diagnosis relies on recognizing the characteristic radiographic and, when available, histopathologic features of usual interstitial pneumonia while excluding other causes of fibrotic interstitial lung disease, and the disease becomes increasingly common with advancing age [1,3]. Recent guidelines increasingly emphasize multidisciplinary evaluation and recognize that characteristic clinical and imaging findings can often establish the diagnosis without surgical biopsy [1]. Clinically, patients most commonly develop worsening breathlessness and persistent dry cough, although the disease often remains unrecognized early and may initially be mistaken for more common cardiopulmonary disorders, contributing to delays in diagnosis [2]. Pathobiologically, IPF is believed to arise when normal repair of the injured alveolar epithelium becomes progressively dysregulated, leading to persistent fibroblast activation, excessive matrix deposition, and progressive disruption of distal lung structure [4]. Emerging evidence further suggests that disease progression reflects coordinated alterations in epithelial repair programs, immune signaling, and stromal function that collectively shape the fibrotic lung microenvironment [4–6]. Single-cell transcriptomic atlases have sharpened this multicellular view by identifying aberrant epithelial states, activated mesenchymal lineages, profibrotic macrophage and immune populations, and altered vascular compartments in IPF lungs [7–9]. Fibroblasts are increasingly viewed not simply as terminal extracellular-matrix producers but as active regulators of tissue repair, local immune tone, and fibrotic persistence in the injured lung [5]. At the same time, the relative contribution of immune and inflammatory processes appears to vary among patients, highlighting the biological heterogeneity that exists within clinically defined IPF [4].

Transcriptomic profiling has become an important tool for studying IPF because it captures the tissue-level consequences of fibrosis and accompanying epithelial, stromal, and immune-associated expression changes. Publicly available RNA-seq datasets now make it possible to study these patterns across larger aggregated sample sets than would be available in most single cohorts [10–14]. However, many existing studies were designed around individual cohorts, anatomical sampling schemes, or focused biological questions. It therefore remains difficult to determine which disease-associated expression programs are reproducible across datasets and which reflect cohort composition, sampling strategy, processing differences, or incomplete metadata.

To address this, we remapped raw sequencing reads from multiple publicly available bulk lung RNA-seq studies and analyzed the resulting count data within a common study-aware framework. This approach reduces processing heterogeneity before biological comparisons are made and allows repeated sampling, study structure, library layout, sex, and age to be handled directly when the necessary metadata are available. The present study was designed to quantify the shared IPF-associated transcriptomic program, test whether disease effects differ by sex, assess whether age modifies disease-associated expression, and summarize the resulting pathway and coexpression structure for biological interpretation.

## 2. Methods

### (a) Study selection

Candidate datasets were identified by searching the Gene Expression Omnibus (GEO) database for idiopathic pulmonary fibrosis RNA-seq studies and manually reviewing study records, associated Sequence Read Archive (SRA) accessions [15], and available sample metadata. The GEO search was last verified on June 5, 2026, using two search terms, “Idiopathic pulmonary fibrosis” and “IPF”, with filters for “Expression profiling by high-throughput sequencing” and “Homo sapiens”. Studies were retained if they represented human bulk lung RNA-seq, included IPF and healthy or nonfibrotic control lung samples, had raw sequencing data available for remapping through SRA, and provided sufficient sample-level metadata for the planned covariate-aware analysis, especially age and sex. Studies were excluded if raw reads were unavailable or not mappable from SRA, if age or sex metadata were unavailable, if the study lacked an appropriate comparison group, or if the data were not compatible with a bulk lung RNA-seq count-based analysis.

The searches identified 355 records. After removing 145 duplicate records between searches, 210 records were screened; 179 were excluded at screening, leaving 31 studies for retrieval and eligibility assessment. Of the 31 studies assessed, 26 were excluded because they did not use bulk RNA-seq (n = 7), lacked raw sequencing data (n = 7), used cultured cells rather than primary lung tissue (n = 6), lacked lung tissue samples (n = 5), or represented a duplicate dataset mapping (n = 1). Five GEO studies met the inclusion criteria and were included in the raw-read reanalysis: GSE138239, GSE184316, GSE199949, GSE213001, and GSE73189 [10–14]. These studies corresponded to 233 samples before metadata filtering: 143 samples from patients with IPF and 90 healthy or nonfibrotic control samples. A study-selection flow diagram documenting this process is included in the accompanying GitHub/Zenodo release.

### (b) Raw sequence processing and count generation

After study selection, SRA run accessions were downloaded with the SRA Toolkit and converted to FASTQ files with fasterq-dump, using split-read output and removal of technical reads where applicable, and read quality was assessed with FastQC before and after read processing [16]. For BioSamples represented by multiple SRA runs within a study, run-level FASTQ files were concatenated by BioSample before alignment.

Reads were processed with Trimmomatic using Illumina adapter clipping and read-length-specific cropping, based on FastQC read quality assessment [17]. Trimmed reads were aligned with STAR using GENCODE v49 annotation, and gene-level counts were generated with STAR -quantMode GeneCounts [18,19], adjusting for single or paired-end libraries per sample as necessary. Strandedness was inferred from the relative total sense and antisense counts.

### (c) Input data and sample harmonization

The resulting bulk lung RNA-seq count matrices were harmonized at the sample-metadata and gene-count levels. For studies in which multiple runs represented the same BioSample, the metadata were collapsed to the BioSample level. Gene-level count matrices were aligned to sample metadata by run accession or merged BioSample identifier, and Ensembl gene identifiers were mapped to gene symbols where available. Samples were retained for analysis if they represented healthy or IPF lung, and had complete information for sex, age, study, library layout, subject sample identifier (for repeated samples), and run accession.

After filtering, 223 samples remained for analysis: 88 healthy and 135 IPF. The filtered dataset comprised 31 healthy females, 57 healthy males, 27 IPF females, and 108 IPF males. The retained studies were GSE138239 (15 samples), GSE184316 (64), GSE199949 (34), GSE213001 (101), and GSE73189 (9). Ten samples were excluded because age was missing. Sixteen subjects contributed repeated samples, and repeated sampling was addressed in the linear model as described below.

(d) Preprocessing and normalization

Gene-level count data were analyzed using `edgeR` and `limma` [20,21]. Low-expression genes were removed with `filterByExpr`, using the four disease-sex groups (healthy female, healthy male, IPF female, and IPF male) as the grouping factor. Library sizes were normalized using trimmed mean of M-values (TMM) [22]. After expression filtering and normalization, 20,310 genes were retained for downstream modeling.

### (e) Linear modeling and differential expression

Differential expression was estimated with a linear modeling framework designed to separate disease effects from major sources of study structure while retaining sex- and age-specific contrasts. Age was standardized to a *z* score across retained samples, and the design matrix was parameterized without an intercept so that the four disease-sex groups were represented directly:

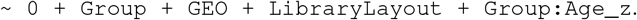

Here, Group denotes the four disease-sex groups: healthy female, healthy male, IPF female, and IPF male. This formulation estimates disease effects at the mean age of the retained cohort, adjusts for study and library layout, and allows the age-expression relationship to differ across disease-sex groups. Sequencing platform was not included as a separate fixed effect because it was completely confounded with GSE184316 and therefore could not be estimated independently of study.

Mean-variance trends were modeled with `voom`, and repeated samples from the same subject were accounted for using `duplicateCorrelation` within the `limma` framework [21,23]. The final model was fit with the subject-level blocking structure, and contrast-level inference used empirical Bayes moderation with robust variance estimation. The primary contrasts estimated IPF-associated expression differences in females, males, and averaged across sex, with additional contrasts evaluating sex differences, disease-by-sex interaction, and disease-associated differences in age slope.

Model outputs were visualized to summarize cohort structure, the dominant disease signal, and overlap between sex-stratified disease contrasts. Principal component analysis was carried out on the 1,000 most variable filtered log counts per million (logCPM) genes as a quality-control visualization. For batch-adjusted visualization outputs, including the disease-gene heatmap, `removeBatchEffect` was used to remove study and library-layout effects, while preserving disease-sex group structure. Volcano plots used FDR *<* 0.05 and absolute log_2_ fold change (| log_2_ FC|) *≥* 1 to highlight differentially expressed genes. Differential-expression tables for all fitted contrasts and visualization outputs are provided with the GitHub/Zenodo analysis release.

### (f) Gene set enrichment analysis

Pathway-level structure was evaluated with preranked gene set enrichment analysis using fgsea [24]. For each contrast, genes were ranked by the moderated *t* statistic, mapped to gene symbols, and collapsed to one representative per symbol by retaining the entry with the strongest absolute statistic. Reactome [25], canonical Kyoto Encyclopedia of Genes and Genomes (KEGG) [26], and Gene Ontology [27] Biological Process (BP), Molecular Function (MF), and Cellular Component (CC) gene sets were obtained from `msigdbr` [28]. KEGG analyses used canonical `C2:CP:KEGG` pathways when available, otherwise `C2:CP:KEGG_LEGACY; KEGG_MEDICUS` terms were not used. Duplicate pathway-gene memberships were removed, and gene sets were restricted to 10-500 genes. Fibrosis-related gene sets were additionally summarized by pathway-name matching for “fibrosis”, “pulmonary fibrosis”, and “interstitial lung”.

### (g) Exploratory network construction

To summarize whether strongly disease-associated genes also showed altered coordination in IPF, we constructed an exploratory differential coexpression network from batch-adjusted voom expression values. Candidate genes were selected from the disease-average results using FDR *<* 0.05 and | log_2_ FC| *≥* 1, then limited to the top 150 genes by significance. Repeated samples were collapsed to subject-level mean expression profiles before network construction. Spearman correlation matrices were estimated separately in healthy and IPF subjects, and differential correlation was defined as *ρ*IPF *− ρ*Healthy. To generate a compact exploratory visualization, gene-pair edges were retained if they showed a large between-condition correlation shift (|*Δρ*| *≥* 0.5) and at least one condition-specific correlation had absolute value *≥* 0.6. Node degree was used as a simple hub summary. This analysis was intended to summarize disease-associated coordination rather than serve as a full module-preservation or weighted gene coexpression network analysis.

### (h) Downstream fixed-signature classifier analysis

To evaluate whether the pooled disease signature retained discriminative value across studies, we performed a downstream fixed-signature IPF-versus-healthy classification analysis. Repeated samples were first aggregated to subject-level units by summing counts across runs within each GEO, subject, condition, and sex combination, yielding 175 units for classification. Gene-level counts were transformed to logCPM with a pseudocount of 0.5. Candidate panels were fixed before model fitting by selecting the top 10, 25, and 50 genes from the disease-average differential expression table among genes meeting FDR *<* 0.05 and | log_2_ FC| *≥* 1.

Cross-study transportability was assessed with leave-one-study-out testing, in which each model was trained on all but one study and evaluated on the held-out study. For each held-out study, an elastic-net logistic regression model with balanced class weights was trained on the remaining studies after standardizing the selected genes within the training set only [29,30]. Regularization was tuned within the training studies by an inner leave-one-study-out search, using balanced accuracy to choose among five values of the scikit-learn penalty parameter *C* (0.01 to 100 on a log scale) and three elastic-net mixing values (*l*_1_ ratio 0.25, 0.5, and 0.75). For each panel, metrics were first calculated within each held-out study and then averaged across studies, including balanced accuracy, receiver operating characteristic area under the curve (ROC AUC), sensitivity, specificity, and precision. Confusion summaries were reported as mean row-normalized held-out-study percentages with study-level standard deviations.

## 3. Results

### (a) Cohort composition and integrated modeling framework

The final integrated analysis retained 223 samples after enforcing complete metadata for disease status, sex, age, study, library layout, and subject identity (Table 1). The modeling strategy was intentionally conservative in two respects: (i) age was modeled linearly within each disease-sex group to maintain an interpretable model that could be estimated reliably across all retained samples, and (ii) sequencing platform was not modeled separately because Ion Torrent data came only from GSE184316, making platform indistinguishable from study. Subject-level blocking was important: the estimated consensus within-subject correlation was 0.522, confirming non-independence among repeated samples.

**Table 1.** Integrated cohort composition after filtering to complete metadata.

| Category | Group | Samples |
| --- | --- | --- |
| Condition | Healthy | 88 |
| Condition | IPF | 135 |
| Sex and condition | Healthy female | 31 |
| Sex and condition | Healthy male | 57 |
| Sex and condition | IPF female | 27 |
| Sex and condition | IPF male | 108 |
| Study | GSE138239 | 15 |
| Study | GSE184316 | 64 |
| Study | GSE199949 | 34 |
| Study | GSE213001 | 101 |
| Study | GSE73189 | 9 |
| Library layout | Paired-end | 31 |
| Library layout | Single-end | 192 |

### (b) A strong cross-study disease signature distinguishes IPF from healthy lung

The integrated disease contrast yielded the dominant signal in the dataset. The average disease comparison identified 8,896 genes at FDR *<* 0.05, including 2,443 genes with FDR *<* 0.05 and | log_2_ FC| *≥* 1 (Table 2). The most significant disease-associated genes included *DIO2, PTGFRN, FNDC1, FHL2, DNAJC22, COL14A1, TSHZ2, MMP16, SSC5D*, and *LTBP1* (Table 3). Many of these genes map to extracellular matrix remodeling, epithelial remodeling, or stromal activation, consistent with a strong IPF remodeling axis. The top-50 disease-gene heatmap showed a coherent separation of healthy and IPF samples across studies after visualization-level batch adjustment (Figure 2A), and the disease-average volcano plot showed that the effect was broad rather than driven by a narrow set of extreme outliers (Figure 2B).

**Table 2.**
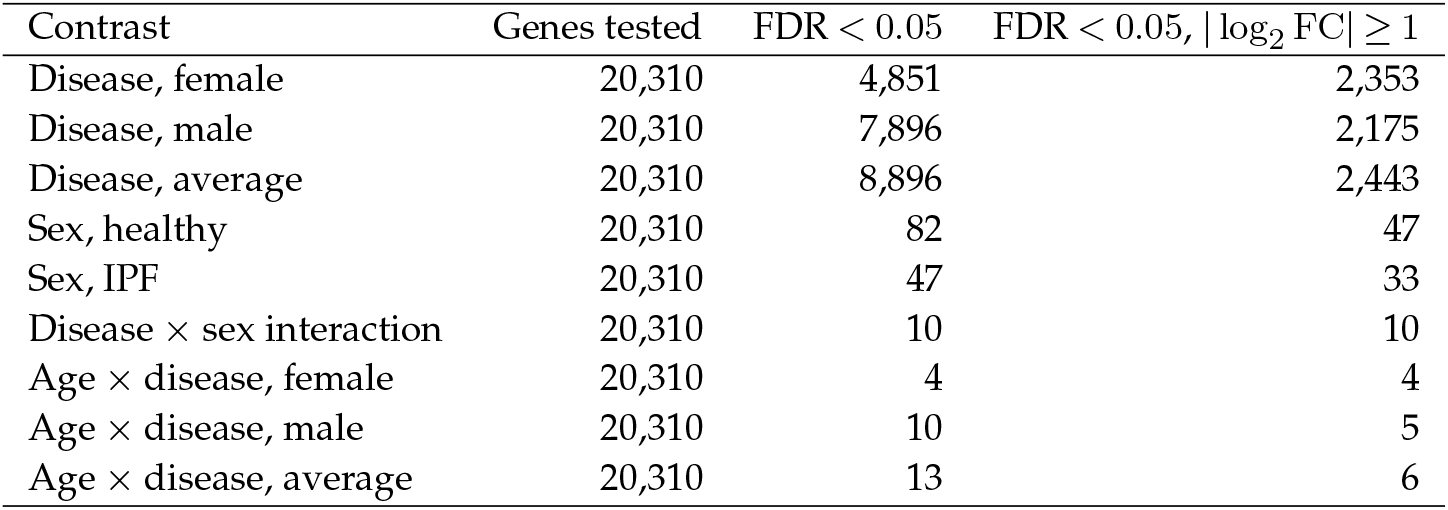
Differential expression summary across all fitted contrasts.

**Table 3.** Top-ranked genes from the disease-average IPF versus healthy contrast (FDR *<* 0.05, | log_2_ FC| *≥* 1).

| Gene | $\log_2 FC$ | FDR |
| --- | --- | --- |
| *DIO2* | 3.790 | $1.50 \times 10^{-29}$ |
| *PTGFRN* | 1.613 | $6.29 \times 10^{-29}$ |
| *FNDC1* | 3.656 | $6.29 \times 10^{-29}$ |
| *FHL2* | 1.932 | $1.89 \times 10^{-28}$ |
| *DNAJC22* | 2.915 | $2.18 \times 10^{-28}$ |
| *COL14A1* | 2.643 | $8.38 \times 10^{-27}$ |
| *TSHZ2* | 1.948 | $1.32 \times 10^{-26}$ |
| *MMP16* | 3.152 | $8.73 \times 10^{-25}$ |
| *SSC5D* | 2.001 | $1.29 \times 10^{-24}$ |
| *LTBP1* | 1.503 | $2.23 \times 10^{-24}$ |

**Figure 1.**
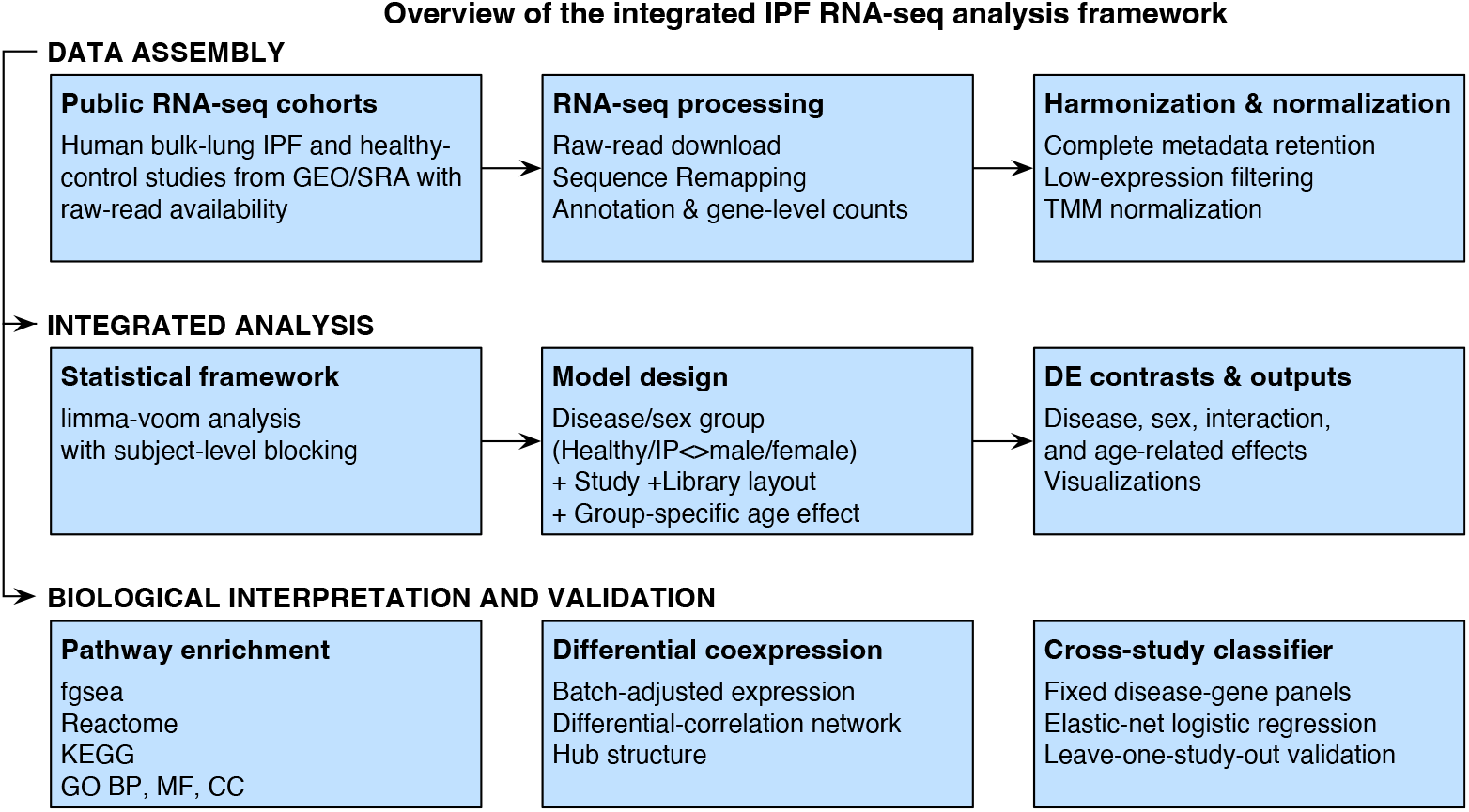
Analysis workflow used to generate the integrated IPF results and downstream fixed-signature classifier reported in this manuscript.

**Figure 2.**
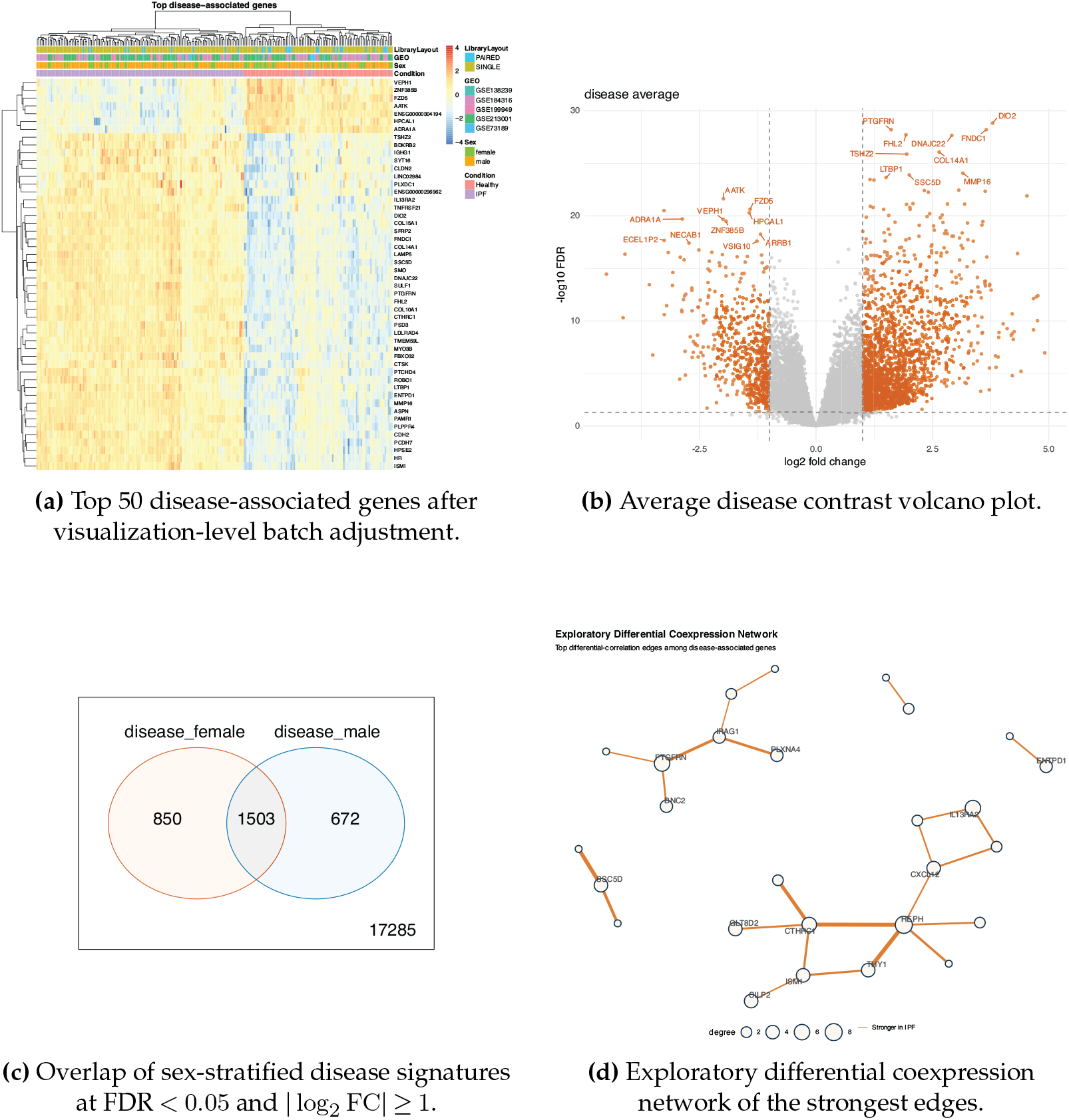
Integrated expression and network signatures of IPF.

### (c) Sensitivity analyses

We performed robustness checks on the main analysis: refitting after the exclusion of the confounded GSE184316 study, refitting without age in the model, and leave-one-study-out disease analyses for each retained GEO study. The first two checks supported the stability of the main disease result. Excluding GSE184316 preserved the disease-average effect almost completely, with Pearson correlation 0.992 against baseline disease log fold-changes and 2,323 shared significant genes at FDR *<* 0.05 and | log_2_ FC| *≥* 1. Removing age also preserved the dominant disease signal, with Pearson correlation 0.974 and 2,219 shared significant genes. These results indicate that the main IPF disease signature is neither driven by the confounded platform/study combination nor dependent on inclusion of the age term.

Leave-one-study-out analyses were also stable for most studies. Omitting GSE138239, GSE199949, or GSE73189 preserved the disease-average signature with Pearson correlations of 0.989, 0.996, and 0.999, respectively, relative to baseline. In contrast, omitting GSE213001 reduced the strength and concordance of the pooled disease result, with Pearson correlation 0.774 and 359 shared significant genes. Additional sex-stratified sensitivity checks indicated that this study contributes disproportionately to the female side of the pooled comparison, whereas the male disease contrast remained more stable. Overall, these robustness analyses support the main disease signal while indicating that subgroup interpretation, especially by sex, should remain cautious. The full sensitivity-analysis results are provided in the GitHub/Zenodo analysis release.

### (d) Fixed disease-gene panels classify IPF across held-out studies

To test whether the pooled disease signature retained discriminative value across cohorts, we performed a downstream subject-level leave-one-study-out classifier analysis using fixed disease panels. The top-50 panel performed best among the prespecified top-10, top-25, and top-50 panels, with macro balanced accuracy 0.940, sensitivity 1.000, specificity 0.880, precision 0.914, and mean ROC AUC 0.950 (Figure 4A). On pooled held-out subject-level predictions, the top-50 panel yielded 105 true positives, 67 true negatives, 3 false positives, and 0 false negatives. Performance was perfect in held-out GSE213001, GSE138239, GSE184316, and GSE199949, whereas the smallest held-out study, GSE73189, showed reduced specificity with 3 of 5 healthy subjects classified as IPF.

**Figure 3.**
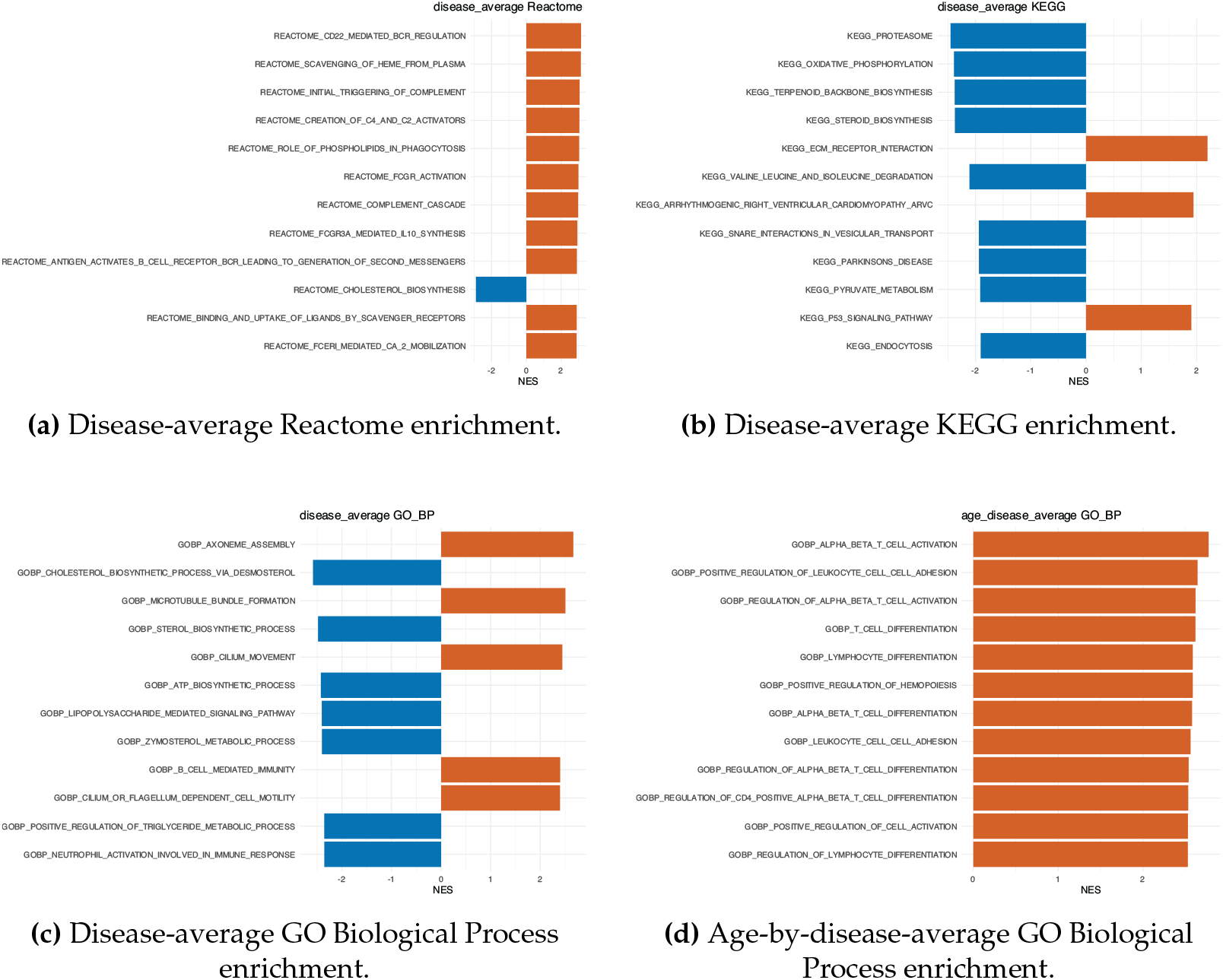
Pathway-level interpretation of the integrated IPF transcriptome.

**Figure 4.**
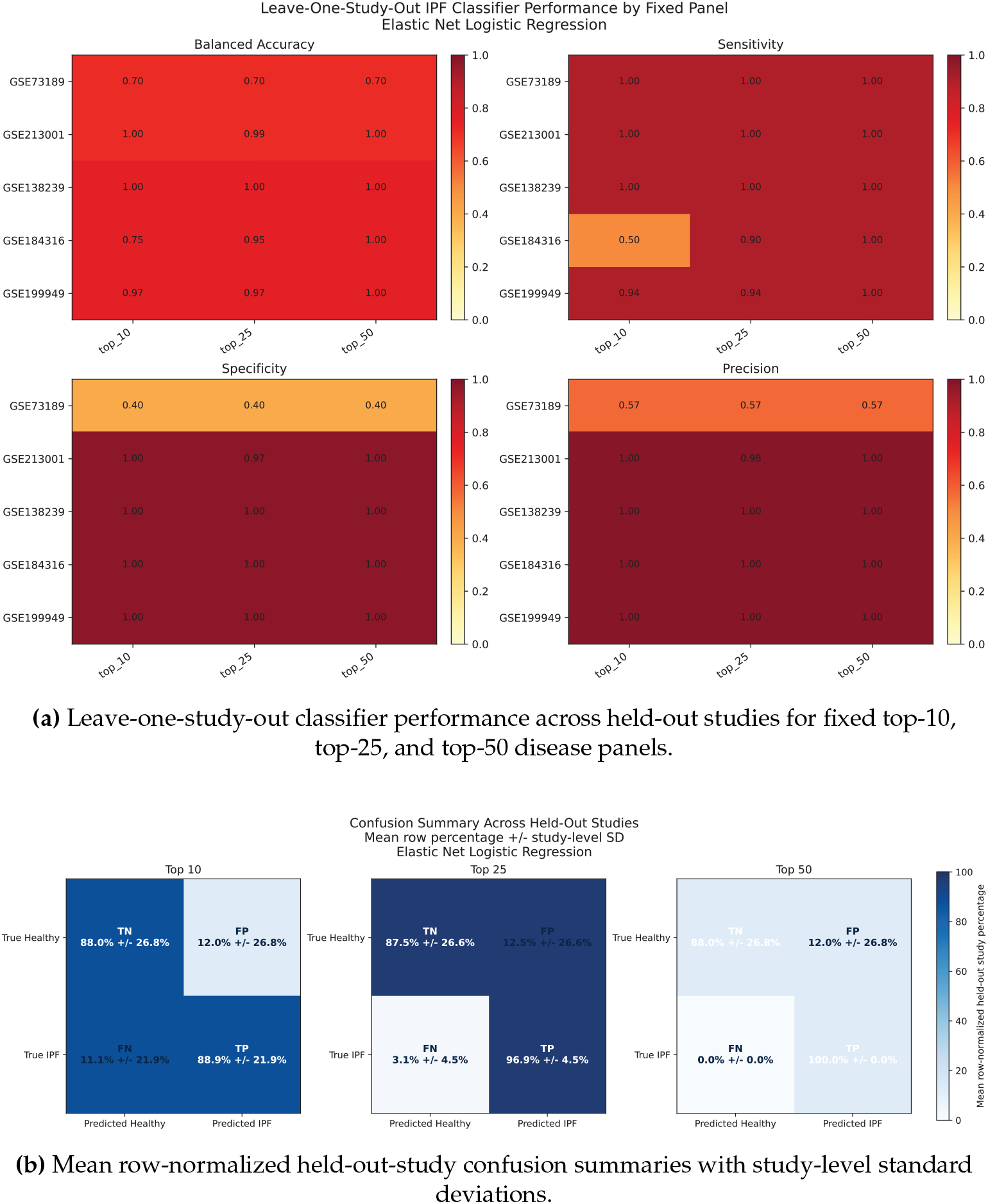
Leave-one-study-out performance of the fixed-signature elastic-net IPF classifier. The top-50 gene panel achieved the best macro balanced accuracy and perfect study-averaged IPF recall, with reduced specificity concentrated in the smallest held-out cohort.

Study-averaged row-normalized confusion summaries therefore showed 100.0% *±* 0.0% true-IPF recall and 88.0% *±* 26.8% true-healthy recall for the top-50 panel (Figure 4B). These results suggest that the dominant pooled IPF signature is portable across most retained cohorts, while also revealing a cohort-specific transportability limit in the smallest study.

### (e) Male and female disease contrasts share a core fibrotic program but are not identical

The disease comparison remained strong in both sexes. In females, 4,851 genes were significant at FDR *<* 0.05, led by *DIO2, FNDC1, IL13RA2, CLDN2, TSHZ2, SYT16, PTCHD4*, and *COL14A1*.

In males, 7,896 genes were significant, led by *FHL2, PTGFRN, DIO2, DNAJC22, MMP16, FNDC1, SULF1, SMO, ISM1*, and other matrix-associated genes. The overlap analysis showed 1,503 genes significant in both sexes at FDR *<* 0.05 and | log_2_ FC| *≥* 1, with 850 female-only and 672 male-only genes (Figure 2C). This is consistent with a large shared fibrotic program plus substantial sex-stratified biology or power differences.

Compared with the sex-stratified disease contrasts, the disease-by-sex interaction results were more focused, identifying only 10 genes passing FDR *<* 0.05. The strongest interaction terms included *CUTALP, CLDN2, ACE2, SYT16*, and several uncharacterized loci. This result suggests that the IPF program is largely shared across sexes, with relatively few genes showing a strong differential disease effect between females and males after adjusting for study and layout.

### (f) Age modifies a smaller but biologically coherent component of the IPF transcriptome

Age-dependent disease effects were much smaller than the primary disease signal. The female age-by-disease contrast yielded 4 genes at FDR *<* 0.05, the male contrast yielded 10, and the average age-by-disease contrast yielded 13. The top annotated age-associated disease genes included *CIC, CNGA1, SNRPF-DT, SYCE1, GPS2P1, RPL29P4, LINC03125*, and *CST7*, alongside several uncharacterized transcripts.

While the age effect was modest at the single-gene level, pathway-level signals were more interpretable and informative. Gene set enrichment for the age-by-disease-average contrast was dominated by adaptive and leukocyte-associated biology, including lymphocyte differentiation, leukocyte cell-cell adhesion, T-cell differentiation, regulation of T-cell activation, positive regulation of leukocyte adhesion, and regulation of hemopoiesis. This pattern suggests that aging in IPF is associated less with a broad shift in canonical matrix genes and more with altered immune cell activation, trafficking, and differentiation states. Age-specific enrichment tables and supplementary age-gene associations are available in the GitHub/Zenodo analysis release.

### (g) Pathway enrichment links IPF to matrix remodeling, humoral immunity, complement, and altered metabolism

Reactome analysis of the disease-average contrast showed strong positive enrichment for scavenging of heme from plasma, CD22-mediated B-cell receptor regulation, phospholipids in phagocytosis, complement cascade, Fc receptor activation, scavenger receptor biology, and extracellular matrix organization (Figure 3A). The same Reactome analysis also showed negative enrichment for translation, cholesterol biosynthesis, and proteasome-associated pathways. KEGG analysis complemented this by showing increased ECM-receptor interaction but relative depletion of oxidative phosphorylation, proteasome-related programs, endocytosis, and cholesterol/mevalonate biosynthesis (Figure 3B). GO Biological Process enrichment additionally highlighted cilium movement, cilium organization, microtubule-based movement, and external encapsulating structure organization (Figure 3C). Overall, these findings indicate that integrated IPF lung tissue is characterized by stromal remodeling and humoral/phagocytic immune signaling, alongside relatively lower representation of metabolic and proteostatic gene programs.

Although both male and female disease contrasts shared the same broad pathway classes, the leading Reactome signals in females emphasized Fc receptor and inflammatory immune modules, whereas the male contrast gave particularly strong enrichment for scavenger receptor uptake, B-cell receptor regulation, complement, and extracellular matrix organization. The similarity between the sex-stratified pathway outputs reinforces the dominant shared disease signal despite incomplete overlap at the single-gene level. Sex-specific pathway enrichment tables are available in the GitHub/Zenodo analysis release.

### (h) Exploratory differential coexpression identifies an IPF-centered remodeling network

The exploratory differential correlation analysis identified an IPF-specific network among top disease genes. The highest-degree nodes were *HEPH, IL13RA2, PTGFRN, CTHRC1, ISM1, CXCL12, GLT8D2, THY1, CILP2*, and *SSC5D*. Several of the strongest edge gains in IPF connected genes including *THY1*-*HEPH, CTHRC1*-*HEPH, SSC5D*-*S100A8, CTHRC1*-*COMP*, and *PTGFRN*-

*IRAG1*. These genes are consistent with activated remodeling, matrix organization, stromal signaling, and altered intergene coordination in IPF and support the interpretation that the integrated IPF signal contains not only changes in mean expression but also reorganization of transcriptional coordination (Figure 2D). Because this network view is threshold-based and exploratory, it should be interpreted as a visualization of differential-correlation structure rather than a formal module-discovery framework.

## 4. Discussion

This integrated analysis identifies a robust cross-study IPF transcriptomic program dominated by matrix remodeling, stromal activation, epithelial remodeling, complement and Fc receptor biology, and strong evidence of humoral immune involvement. The top-ranked genes and enriched pathways are consistent with a tissue-level fibrotic microenvironment composed of activated mesenchymal compartments, altered immune cell composition, and remodeling of extracellular architecture, which fits the broader pathogenic framework described for IPF [2, 4]. The simultaneous enrichment of extracellular matrix organization and B-cell/complement pathways suggests that, in these bulk lung profiles, fibrosis and immune activation are tightly coupled rather than separable secondary effects. The accompanying epithelial-remodeling and cilium-associated signals are likewise consistent with newer multicellular models of IPF that emphasize dysregulated alveolar epithelial repair, bronchiolization, abnormal epithelial-cell states, and persistent profibrotic cross-talk among epithelial, mesenchymal, immune, and endothelial compartments [4,7–9]. These single-cell atlases provide cellular context for the bulk signature, while the present analysis tests whether the aggregate tissue-level program is reproducible across cohorts. This pattern is also compatible with the emerging view that fibroblasts help organize immune-cell recruitment, repair failure, and matrix persistence in fibrosing lung disease rather than acting only as passive structural effectors [5].

Several aspects of the present results are also concordant with prior transcriptomic studies represented in the integrated cohort. The persistence of strong extracellular-matrix and remodeling signals aligns with prior observations from the viral-assessment cohort, the central-versus-peripheral IPF cohort, and the apex-versus-base cohort, even though those studies were designed around different primary questions [10,12,13]. Yin and colleagues, analyzing the viral-assessment cohort by RNA-seq, detected only sporadic low-level viral reads and found no statistically supported enrichment of any virus in IPF relative to control lungs. In that context, the dominant cross-study signal recovered here is more consistent with host remodeling and immune-state dysregulation than with a broadly shared active viral RNA signature [10]. In particular, Huang and colleagues reported that relatively central lung regions in IPF, despite being less overtly remodeled histologically than peripheral fibrotic zones, already display transcriptional features linked to activated myofibroblasts and idiopathic pulmonary fibrosis signaling pathways; the present pooled analysis is consistent with that view and extends it by showing that remodeling-associated disease programs remain prominent after harmonizing across multiple cohorts rather than within a single spatially sampled study [12]. Jaffar and colleagues likewise showed that MMP7 expression increases in more fibrotic basal IPF tissue relative to apical regions, supporting the broader concept that regional progression of fibrosis is accompanied by strengthening matrix-remodeling programs; our pooled analysis is concordant with that framework, although the integrated disease signal is not limited to any single remodeling mediator [13]. Yu and colleagues further implicated reduced BMP3 expression as part of a permissive profibrotic program in idiopathic interstitial pneumonias, which is directionally consistent with our broader observation that the pooled IPF transcriptome is dominated by matrix-promoting and stromal-activation signals rather than by antifibrotic restraint [14]. The current pooled analysis extends those observations by estimating a study-adjusted disease effect across all retained cohorts within a common analytical framework that explicitly accounts for repeated sampling, age, and sex.

The sex-stratified comparisons indicate that most disease biology is shared, but they also show that the shared set is incomplete. Female IPF retained strong signals in *IL13RA2, CLDN2, SYT16*, and immunoglobulin-related genes, whereas male IPF more strongly highlighted *FHL2, MMP16, SULF1, SMO, ISM1*, and *ASPN*. Because the formal interaction signal was small, these sex differences are better interpreted as secondary quantitative shifts superimposed on a largely common disease architecture rather than as evidence for two distinct molecular diseases.

At the single-gene level, age-by-disease effects were also limited, but at the pathway level aging clearly tracked immune activation, cell adhesion, and T-cell differentiation programs. This suggests that age in IPF may act more through coordinated changes in immune state and tissue composition than through large age-specific shifts in a small set of canonical fibrosis genes. These observations suggest that aging in IPF influences the immune environment within an already remodeling and fibrotic lung, rather than pointing to a distinct inflammatory process [3].

The sensitivity analyses provide context for the robustness of the main disease results. The pooled signature was stable after excluding the confounded GSE184316 study and after removing age from the model, but the leave-one-study-out analysis showed that GSE213001 contributes substantially to the pooled estimate. This pattern is compatible with a consistent shared disease signal that is strengthened by the most informative cohort rather than a simple technical artifact, but it also means that our conclusions should rest on the most consistent genes and pathways across studies and that sex-specific claims should be more cautious.

A strength of this study is that the retained cohorts were reprocessed from raw sequencing reads rather than relying on heterogeneous study-provided count matrices, improving comparability of gene-level quantification across datasets. Unlike conventional transcriptomic meta-analyses that combine study-level results after independent processing, the present approach enabled all retained samples to be analyzed within a common statistical framework while explicitly accounting for study structure, repeated sampling, age, and sex. This design strengthens confidence that the dominant matrix-remodeling and immune-associated signals identified here reflect reproducible features of IPF biology rather than artifacts of individual analytical pipelines. However, raw-read reprocessing does not remove differences in cohort design, tissue sampling, library construction, sequencing depth, or metadata completeness, so study-aware modeling and sensitivity analyses remain necessary.

The downstream fixed-signature classifier supports that same interpretation from a predictive angle. A top-50 gene panel taken directly from the pooled disease signature classified held-out cohorts with high macro balanced accuracy and no false negatives in pooled subject-level testing, which suggests that the dominant integrated IPF program is not merely cohort-specific noise. This classifier result is best interpreted as evidence that the pooled host-expression signature retains cross-cohort transportability, rather than as a clinical diagnostic model. However, specificity dropped in GSE73189, indicating that transportability is not uniform across all retained cohorts and that even a biologically anchored signature can misclassify healthy subjects from a small study whose expression profiles sit near the IPF decision boundary.

The exploratory coexpression results further support disease-associated rewiring. Hub genes such as *HEPH, IL13RA2, PTGFRN, CTHRC1, ISM1, CXCL12, THY1*, and *CILP2* point to an interconnected remodeling and stromal-signaling module that is strengthened in IPF. Rather than a full validated network, the limited threshold-based network identifies candidate rewiring nodes that may be investigated further in future studies to provide more mechanistic insight into transcriptional regulation in IPF.

Overall, our results suggest that previous transcriptomic findings from studies focused on viral assessment, spatial heterogeneity, regional disease severity, and antifibrotic regulation converge on a common disease-associated program characterized by extracellular-matrix remodeling, epithelial dysfunction, and coordinated immune activation. Thus, by harmonizing raw RNA-seq data across independent patient cohorts and modeling key study and demographic factors directly, our study has identified concordant disease-associated signals that are less likely to reflect single-dataset artifacts. Our findings point to a convergent matrix-immune IPF program that likely represents fundamental disease mechanisms and may provide focus for developing effective treatments across heterogeneous IPF patient populations.

## 5. Limitations

Our study has several constraints. First, the metadata completeness limited our cohort (both in study selection, as well as within the selected studies), as we had to exclude samples missing age annotations. Additionally, disease severity measures (such as spirometry, fibrosis extent, disease duration, or lung location) were not uniformly available across the included studies and could not be uniformly and consistently incorporated into the analysis, which limits discerning the effect of disease severity and tissue/spatial heterogeneity on IPF differential gene expression.

There is also confounding, beyond experimental approaches, including sequencing platforms. For example, for study GSE184316, the unique Ion Torrent platform could not be disentangled from study effects, precluding independent platform adjustment. In terms of modeling, we note that our age modeling used linear terms within each disease-sex group to maintain interpretability, but this may not capture nonlinear age-expression relationships that a spline-based analysis could reveal (which would be more robust with denser data). Furthermore, the aggregate disease signal is naturally influenced by GSE213001, which was the largest study included, and hence we must be cautious in our pattern interpretations, particularly for sex-stratified results.

The exploratory differential coexpression network is threshold-based and does not represent a complete regulatory network; rather, it identifies candidate nodes with rewired connections across differentially expressed genes for further investigation. Similarly, the classifier analysis was designed to evaluate cross-study applicability of a fixed gene-expression-based signature.

Finally, our analysis is based on bulk RNA-seq data, meaning all expression measurements reflect aggregate cell signals rather than resolving cellular subpopulation behavior. Single-cell sequencing and spatial transcriptomics can provide cellular-level resolution, but bulk RNA-seq remains valuable for detecting robust tissue-level differential expression in limited, heterogeneous cohorts such as IPF.

## Data Accessibility

The original raw sequencing data are publicly available through GEO/SRA under accessions GSE138239, GSE184316, GSE199949, GSE213001, and GSE73189. All associated analysis scripts, harmonized inputs, result tables, and figure outputs, including the figures referenced in the manuscript, have been deposited on GitHub and archived as a Zenodo release (DOI: 10.5281/zenodo.20830716; URL: https://github.com/gmiaslab/IPF_Analysis).

## Use of AI disclosure

During preparation of the computational workflows, the authors used AI-assisted coding tools within Visual Studio Code for routine function completion, code debugging, and formatting cleanup. All analytical workflows, study design decisions, quality checks, results validation, and scientific interpretations were performed and verified by the authors.

## Author Contributions

S.N. performed data curation, formal analysis, visualization, and contributed to writing the original draft. J.H. contributed to methodology, and manuscript review and editing. G.I.M. conceived and supervised the study, contributed to methodology and formal analysis, interpreted the results, and contributed to writing, review, and editing. All authors reviewed and approved the final manuscript.

